# Network-guided supervised learning on gene expression using a graph convolutional neural network

**DOI:** 10.1101/2021.12.27.474240

**Authors:** Hatairat Yingtaweesittikul, Chayaporn Suphavilai

## Abstract

**Background:** Transcriptomic profiles have become crucial information in understanding diseases and improving treatments. While dysregulated gene sets are identified via pathway analysis, various machine learning models have been proposed for predicting phenotypes such as disease type and drug response based on gene expression patterns. However, these models still lack interpretability, as well as the ability to integrate prior knowledge from a protein-protein interaction network.

**Results:** We propose *Grandline*, a graph convolutional neural network that can integrate gene expression data and structure of the protein interaction network to predict a specific phenotype. Transforming the interaction network into a spectral domain enables convolution of neighbouring genes and pinpointing high-impact subnetworks, which allow better interpretability of deep learning models. Grandline achieves high phenotype prediction accuracy (67-85% in 8 use cases), comparable to state-of-the-art machine learning models while requiring a smaller number of parameters, allowing it to learn complex but interpretable gene expression patterns from biological datasets.

**Conclusion:** To improve the interpretability of phenotype prediction based on gene expression patterns, we developed Grandline using graph convolutional neural network technique to integrate protein interaction information. We focus on improving the ability to learn nonlinear relationships between gene expression patterns and a given phenotype and incorporation of prior knowledge, which are the main challenges of machine learning models for biological datasets. The graph convolution allows us to aggregate information from relevant genes and reduces the number of trainable parameters, facilitating model training for a small-sized biological dataset.

## Background

Transcriptomic profiles have become key information in understanding diseases and improving treatments [1–3], where high throughput RNA sequencing (RNA-seq) is widely used for quantifying the expression of genes in each patient [4, 5]. One common analysis is to identify differentially expressed genes (DEGs) [6–8], as well as dysregulated pathways via pathway analysis [9, 10], to pinpoint the disrupted biological process and understand disease mechanisms. While identifying DEGs requires phenotypic information of samples, gene expression clustering methods [11] search for common patterns to stratify patients and discover disease subtypes [12, 13]. Both DEGs and clustering methods are useful tools for understanding disease mechanisms, but they could not be directly used for predicting a phenotype of an unseen patient based on gene expression.

Various machine learning models have been proposed for predicting phenotype based on gene expression, ranging from traditional machine learning models [14, 15] to deep learning models [16–18]. The models learn transcriptomic patterns from a large sample cohort to predict phenotypes such as disease status [19, 20] and drug response in unseen samples [21, 22]. However, most standard machine learning models [23–25] do not allow us to incorporate the relationship between genes from a protein-protein interaction (PPI) network [26].

Deep learning models have recently been used for predicting phenotypes based on gene expression profiles [16, 27]. Despite the ability to capture complex patterns, many models still lack transparency [28, 29]. In a standard deep neural network (DNN), a fully connected layer captures a linear combination of all genes but could not utilise protein interaction information from PPI networks. In contrast, a convolutional neural network (CNN), which is widely used in the image processing field [30, 31], contains a convolutional layer that can combine nearby pixels (or related features). Based on the convolutional concepts, Kipf and Welling have proposed a graph convolutional neural network (GCN) that transforms a graph onto a spectral domain, enabling convolution of neighbouring nodes with respect to a given graph structure [32].

Recently, few studies have incorporated PPI network and gene expression. Nguyen *et al*. apply GCN to capture the molecular structure of drugs [33], whereas other studies incorporate PPI networks in the preprocessing steps to construct a feature matrix [34, 35]. However, applications of GCN for integrating gene expression and PPI network to predict a phenotype of an unseen sample has never been explored.

## Results

### Grandline framework

We proposed Grandline, a graph convolutional neural network (GCN) model that transforms PPI network into a spectral domain to enable the convolution of expression values among neighbouring genes. The model integrated gene expression profiles with the structure of PPI network to predict a phenotype of unseen samples (**Figure 1**). By adapting Grad-CAM technique [36], Grandline could also identify subnetworks that play critical roles in predicting a given phenotype, improving interpretability of the model.

**Fig. 1.**
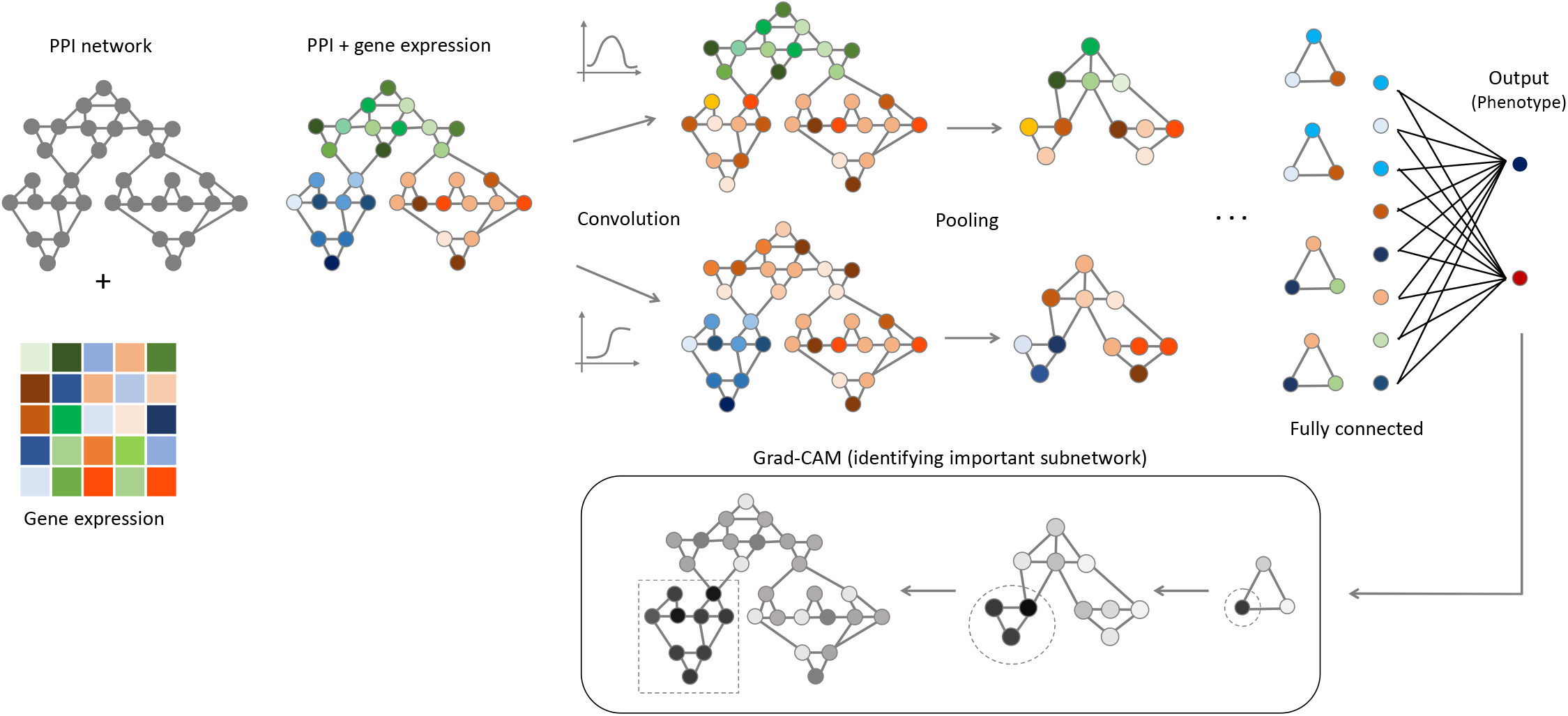
Grandline framework. Gene expression values are overlaid onto a protein-protein interaction network (PPI network). Grandline consists of multiple blocks for convolution and pooling layer. The convolution step is based on a convolutional filter defined on a spectral domain, and the pooling step is based on graph coarsening. After the last pooling step, feature maps are flattened, and multiple fully connected layers are applied for predicting a given phenotype. Finally, Grad-CAM technique is adapted to identify subnetworks that are important for the phenotype prediction of each sample.

Grandline integrates PPI network by considering the network as an undirected graph and gene expression values as node signals. Similar to a standard conventional neural network (CNN) models, the model consists of multiple blocks for convolution and pooling layer. To enable convolution layer that can aggregates expression levels of multiple genes, we defined a spectral graph convolution on the Fourier domain [32, 37] and then defined a convolutional filter based on Chebychev polynomial [38] (**Methods**). For the pooling operation, we applied a graph coarsening method [37] to combine the output of the convolutional layer. Fully connected layers were then used for predicting a given phenotype.

Drawing on the interpretability, Grandline could identify subnetworks that are important for the phenotype prediction using Grad-CAM technique [36] (**Methods**). Originally, this technique is used for overlaying a heatmap onto the input image of a CNN model, highlighting an important region of the image. By adapting Grad-CAM technique to the GCN model, Grandline pinpointed biological subnetworks on a given PPI network that play an important role in predicting a curtain phenotype. This feature allows us to understand the underlying mechanisms of disease and drug response.

### Grandline for supervised-learning and subnetworks identification

Synthetic data were used for evaluating prediction accuracy and the ability to identify correct predefined subnetworks that are related to the target classes. We evaluated the predictive performance and the ability to identify subnetworks on two simulated linear (SIM-L) and nonlinear (SIM-NL) datasets that consist of 5,000 nodes and 200 samples, where we predefined subnetworks that correlate with class labels (**Methods**). Six state-of-the-art methods (KNN, ElasticNet, NaïveBayes, SVM-Linear, SVM-RBF and DNN) for predicting phenotypes based on gene expression patterns were also evaluated.

For the linear dataset, all models achieve > 99% accuracy based on cross-validation, but only Grandline and SVM-Linear correctly pinpointed 6 predefined subnetworks (C3, 6, 9, 13, 16, 19) (**Figure 2A, Supplementary figure 1A**), highlight interpretability of the model. In SVM-Linear, the coefficients in the primal problem can infer feature importance based on its support vectors. While ElasticNet logistic regression model returned feature coefficients, the coefficients only highlight few genes within the predefined subnetworks (**Supplementary figure 1B**). For KNN, NaïveBayes and SVM-RBF, features important could not be directly inferred based on the output models.

**Fig. 2.**
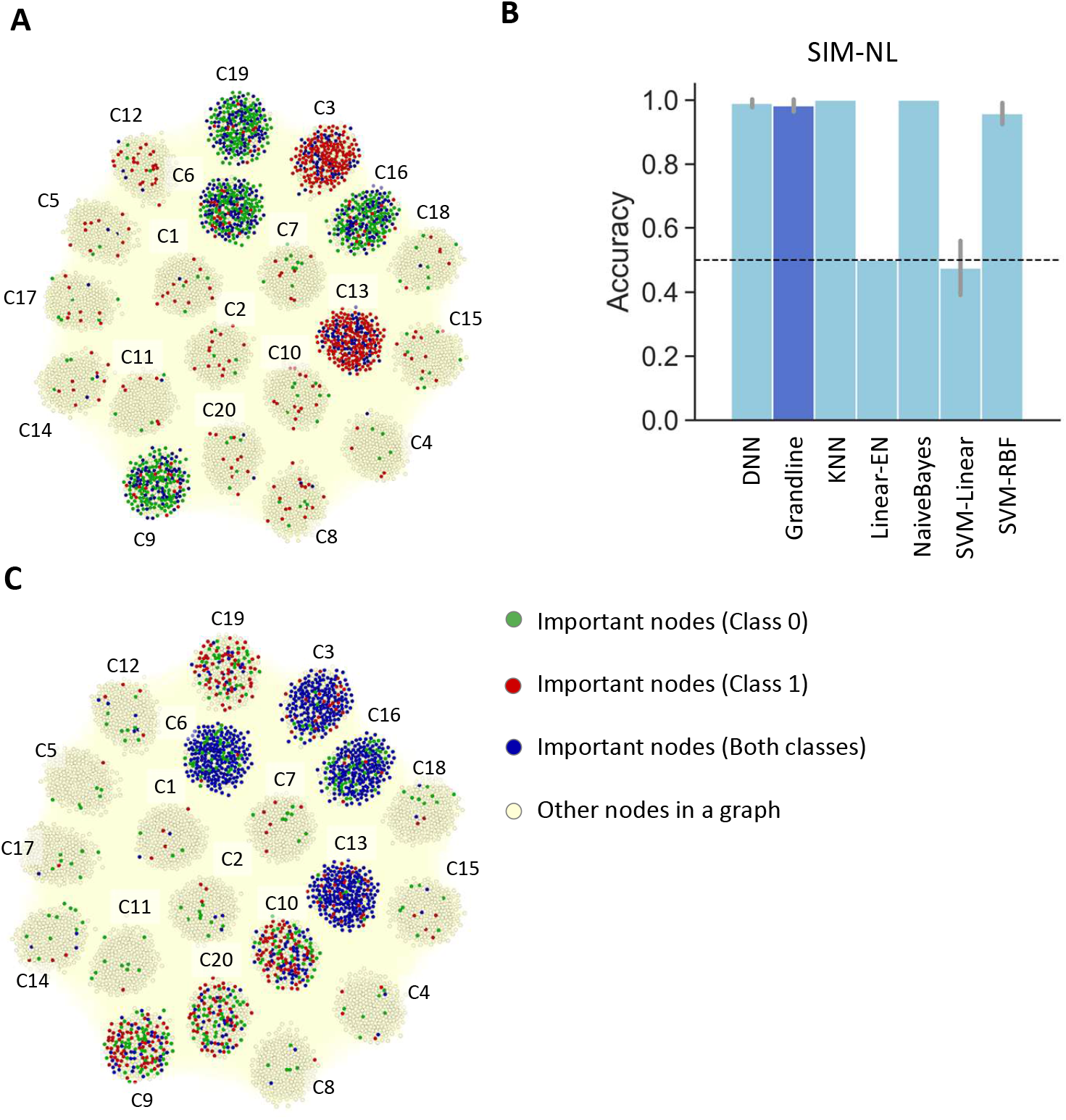
Model performance and interpretability. (A) Important subnetworks (C3, 6, 9, 13, 16, 19) were correctly identified by Grandline in a simulated linear (SIM-L) dataset. For node colors, green, red, blue represents nodes that are important for predicting class 0, class 1, and both classes, respectively. (B) Comparison of average accuracy between Grandline (dark blue) and six state-of-the-art models (light blue). A grey line represents a standard deviation of the accuracy across multiple folds. (C) Correct important subnetworks (C3, 6, 9, 10, 13, 16, 19, 20) identified by Grandline in a simulated nonlinear (SIM-NL) dataset.

Drawing on the nonlinear dataset, 5 methods (KNN, NB, SVM-RBF, DNN, and Grandline) successfully predicted class labels (accuracy > 0.96). As expected, the linear models failed to capture the nonlinear relationship (accuracy ≤0.5) (**Figure 2B**). Amongst the models with high predictive performance, only Grandline could identify all 8 predefined subnetworks (**Figure 2C**). For DNN, the integrated gradients method was applied to assign importance values to each feature based on the gradients of the prediction with respect to the input [39]. While all 6 subnetworks were correctly identified in SIM-L dataset, the integrated gradients method could identify only 6 out of 8 correct subnetworks for SIM-NL dataset (**Supplementary figure 1C-D**). The KNN, NB and SVM-RBF models do not provide feature importance.

Performance evaluation on the simulated datasets demonstrated that Grandline could correctly predict class labels in both linear and nonlinear scenarios, as well as pinpoint the predefined subnetworks. Our results highlight that Grandline could integrate PPI network information with a supervised classification model, enabling us to identify important subnetworks (i.e., pathways or gene sets) that play an important role in phenotype prediction.

### Phenotype prediction and subnetworks identification

As a case study, we obtained two datasets of major depression disease (MDD1 and MDD2) consisting of 157 (79 healthy vs 78 patient) and 59 (29 healthy vs 30 patient) samples [40, 41], respectively (**Methods**). Grandline showed the highest accuracy in both datasets (76.67% and 78.33%) and was notably better than conventional machine learning models, except for SVM-RBF in MDD1 dataset (**Figure 3A, Supplementary figure 2**). These results suggest that deep learning models could capture complex gene expression patterns that distinguish healthy and disease cases. Furthermore, by inspecting the number of parameters used in the DNN model and Grandline for both MDD1 and MDD2 datasets, we observed that the convolutional technique could reduce the number of trainable parameters by 97% and 91% (97,927 vs 3,231,858 and 95,672 vs 1,034,194 parameters), suggesting that Grandline could be more suitable for a small-size data.

**Fig. 3.**
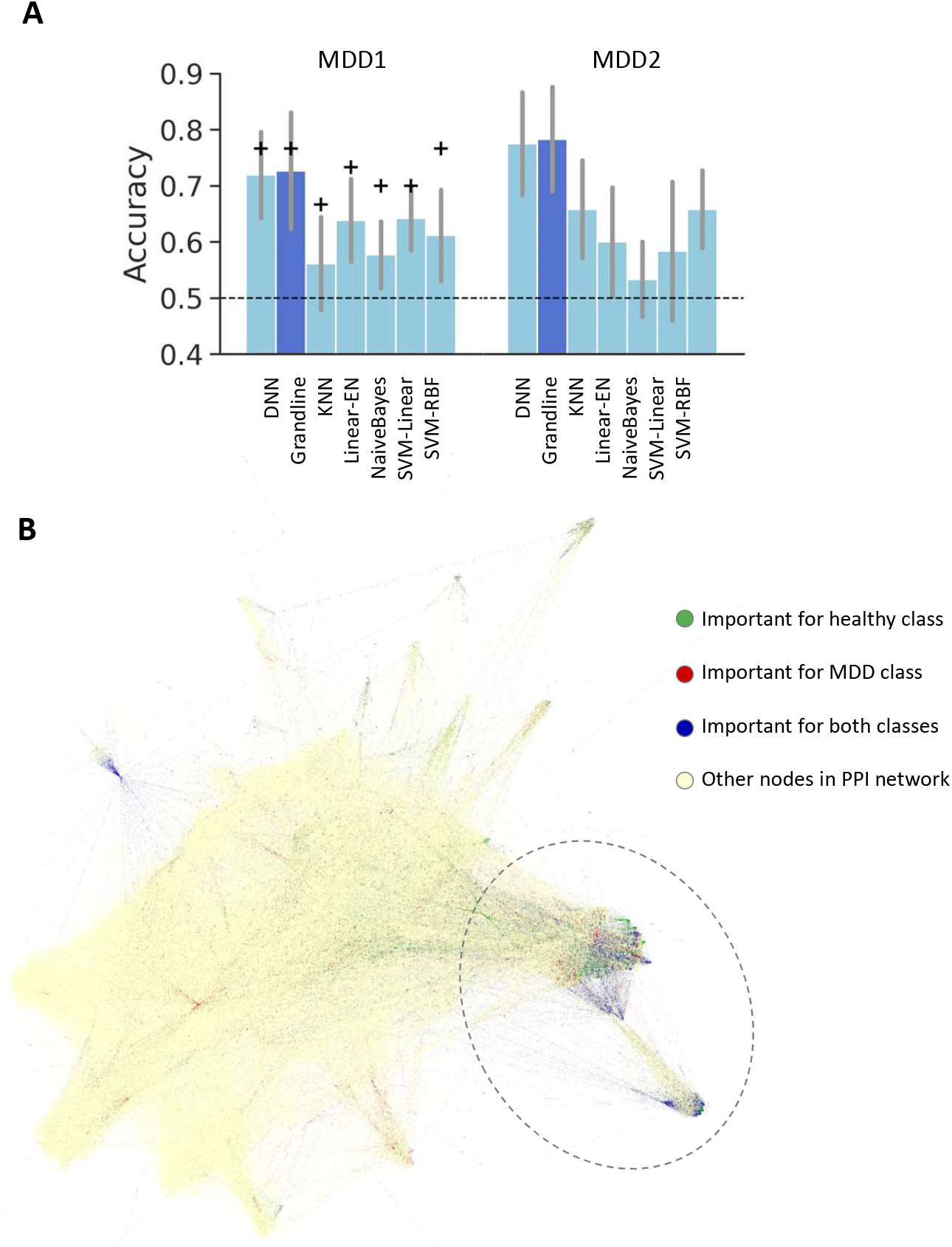
Predictive performance and identified subnetworks for MDD datasets. (A) Comparison of average accuracy between Grandline (dark blue) and six state-of-the-art models (light blue) for MDD1 and MDD2 datasets. A grey line represents a standard deviation of the accuracy across multiple folds (cross-validation), and a marker + represents the accuracy of the test set. (B) Visualisation of PPI network. Grandline identified important subnetworks, which could distinguish healthy and MDD samples for MDD2 dataset.

By adapting Grad-CAM technique used in CNN onto the spectral domain in GCN (**Methods**), Grandline could identify multiple important subnetworks in both datasets. The top 313 and 807 important genes (top 5% highest importance values) were identified for MDD1 and MDD2 datasets, respectively. From network visualisation, we found two subnetworks for MDD2 (at the bottom right corner of the network; **Figure 3B**) that play an important role in the phenotype prediction, as well as important subnetworks for MDD1 (**Supplementary figure 3**). To investigate the important genes identified in the subnetworks, we performed gene set enrichment analysis using DAVID (**Supplementary file 1**). Several pathways that are known to be related to MDD were identified, including Protein processing in the endoplasmic reticulum [42], Neuroactive ligand and Cytokine-cytokine receptor interactions [43], as well as several signaling pathways such as NF-kappa B [44], Chemokine, and Jak-STAT [45, 46]. We also observed that the important genes were enriched in behavioural pathways, including Circadian entrainment [47] and Alcoholism [48]. We note that the numbers of available genes are largely different between MDD1 and MDD2 datasets, leading to different predicted subnetworks. Interestingly, the important genes in both MDD1 and MDD2 were enriched in Salmonella infection, in line with previous studies that reported a link between depression and infection caused by Salmonella [49, 50].

To compare against the state-of-the-art pathway analysis tool, we performed independent GSEA analyses [9] to identify enriched gene sets (**Supplementary file 2**). While GSEA could not identify significantly differentiated gene sets for MDD1, we only observed a small number of overlapping genes (Jaccard index = 0.06) when comparing 503 genes from the top 10 pathways of GSEA result with 1,305 important genes in identified subnetworks of MDD2. The small overlap suggests that our subnetworks could provide additional information for phenotype prediction and understanding disease mechanisms. Taken together, these results highlight the utilities of Grandline to simultaneously predict phenotype and pinpoint important subnetworks based on transcriptomic profiles and PPI networks in real-world datasets.

### Predicting cancer drug response and subnetwork biomarkers

To predict cancer drug response and identify subnetwork biomarkers based on gene expression and PPI network, we obtained gene expression and drug response information of 5 anticancer drugs (525-563 cell lines tested for each drug), including Methotrexate, Oligomycin, Paclitaxel, SB743921 and Vincristine (**Methods**) [21]. In terms of predictive performance, Grandline was among the best for all drugs (accuracy = 67.27%-85.45%) and showed the best performance in Paclitaxel and Vincristine with accuracy 84.90% and 81.25%, highlighting the predictive power of the model (**Figure 4A**; **Supplementary figure 4-5**). Compared to DNN, Grandline could reduce the number of parameters by 81.95% (1,566,664 vs 8,686,832) for SB743921, 63.38% (1,566,694 vs 4,278,322) for Paclitaxel, 52.20% (1,880,119 vs 4,292,770) for Oligomycin, 26.76% (3,133,261 vs 4,278,322) for Methotrexate, and 12.10% (470,066 vs 534,834) for Vincristine. These results suggested that Grandline could learn gene expression patterns on the network to predict drug response using a smaller number of parameters.

**Fig. 4.**
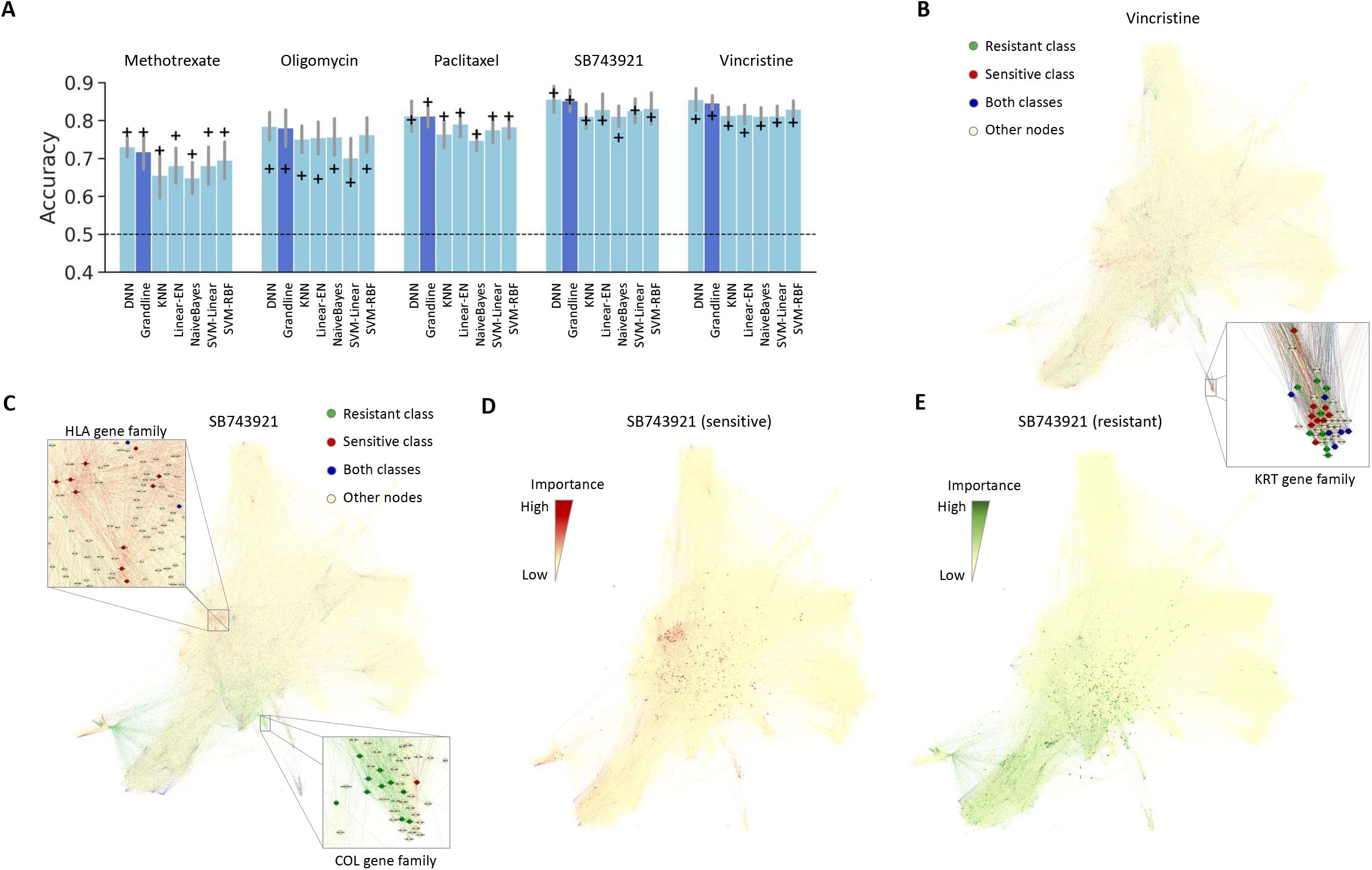
Predictive performance and subnetwork biomarkers of cancer drug response prediction. (A) Comparison of average accuracy between Grandline (dark blue) and six state-of-the-art models (light blue) of the unseen samples across 5 cancer drugs. A grey line represents a standard deviation of the accuracy across multiple folds (cross-validation), and a marker + represents the accuracy of the test set. (B) Important subnetworks identified by Grandline for resistant and sensitive classes of Vincristine. A highlighted subnetwork consists of several *KRT* genes identified for both sensitive and resistant classes. (C) Important genes for resistant and sensitive classes of SB743921, where sensitive (red) and resistant (green) subnetworks appear in different network regions. (D-E) Grandline identifies sample-specific important subnetworks for each cell line that is sensitive and resistant to SB743921.

Drawing on the interpretability of the model through important subnetworks identified by Grandline, we investigated biomarker subnetworks for Vincristine and SB743921, for which Grandline showed the highest accuracy (81.25% and 85.45%). By visualising the PPI network with node importance for different cancer drugs, subnetworks associated with drug response mechanisms were pinpointed. For Vincristine, a subnetwork consisting of *KRT* gene family was identified, in line with the finding that aberrant keratin expression confers drug response phenotypes [51] (**Figure 4B**). For SB743921, which is a chemical inhibitor for kinesin spindle protein that is essential for cell-cycle progression [52], we observed that essential genes for sensitive and resistant classes were showed up in distinct regions of the PPI network (**Figure 4C**). We highlighted human leukocyte antigen (*HLA*) family identified as an important subnetwork for the sensitive class. This result aligned with the previous finding that the kinesin superfamily proteins could bind with human leukocyte antigen molecules and can potentially be used for immunotherapy [53]. Another subnetwork identified for the resistant class is a collagen (*COL*) family, which is upregulated in M phase of the cell cycle, where the kinesin motor protein is also induced [54]. We note that 22% of important genes identified by Grandline was identified by DNN using the integrated gradients methods (**Supplementary figure 6**); however, only 3 (HLA-DOA, HLA-DQB2, HLA-DRB1) and 1 (COL6A1) genes from HLA and COL gene families were detected.

One advantage of Grad-CAM is the ability to identify sample-specific important features. We considered different cell lines, JM1 and SUIT2, for which the model correctly predicted sensitive and resistant responses with the highest probabilities. Distinct subnetworks have been identified for both cell lines (**Figure 4D-E**, red = sensitive and green = resistant), suggesting the unique ability of Grandline to pinpoint sample-specific subnetworks contributing to a specific phenotype. In conclusion, we demonstrated that Grandline could predict phenotype (disease status and drug response) and identify subnetwork biomarkers based on the integration of transcriptomic profiles and PPI network, enabling future applications for studying disease mechanisms and personalised medicine.

## Discussion

Several machine learning models have been proposed for predicting a specific phenotype based on gene expression profile, but only a few models have incorporated the prior knowledge from a protein-protein interaction network. These models usually incorporate the network in feature selection steps [34, 35], whereas direct integration of the interaction network and gene expression to predict a specific phenotype has not been explored. Here we propose a new deep learning framework, Grandline, that can predict a phenotype based on the integration of gene expression profile and protein-protein interaction (PPI) network, resulting in the ability to pinpoint important subnetworks and capture the complex nonlinear relationship between the patterns of gene expression and target phenotype (**Figure 1**). By applying the model on two simulated datasets, as well as multiple real-world datasets (major depressive disorder and drug response on cancer cell lines), we found that Grandline could identify relevant subnetworks without sacrificing predictive performance (**Figure 2-4**).

Although a deep learning model can learn complex patterns, it is usually considered as a ‘black box’ due to its high complexity and lack of explanation/reasoning for the prediction [28]. Recently, several studies have focused on the interpretability of deep learning models [55, 56], where the majority of the methods are designed for DNN and a fewer number for GCN. We note that some recent methods such as GNNExplainer that support convolutional layers are typically designed for the scenario where each node represents a sample/patient (a graph consists of multiple samples, and each node contains multiple signals or features). This scenario is different from our set-up, where each node represents a gene and contains a gene expression value.

We adopted the existing visualisation technique, Grad-CAM [36], into a graph convolutional neural network to identify important subnetworks, which are hard to identify based on existing machine learning models. For instance, a logistic regression model such as ElasticNet would not be able to identify a set of correlated genes within a subnetwork. Besides, using convolutional layers provides advantages over fully connected layers in the standard deep neural network. The convolutional layers allow us to consider multiple relevant genes simultaneously and reduce the number of trainable parameters, facilitating model training for a small-sized biological dataset.

Despite different types of graph convolutional neural networks (GCN) [57], existing models based on biological networks either focus on predicting expression values of neighbouring genes [58], prioritising disease-associated genes [59, 60] or capturing drug structure [33]. Several methods have also been proposed for interpreting GCN models [61, 62]. These GCN-based models are designed for either predicting node label, node interaction or similarity between different graphs. In contrast, Grandline relies on a global network structure but different node signals (gene expression) across samples, enabling the model to simultaneously predict a phenotype and identify distinct important subnetworks for each sample. However, some caveats of this strategy include the fact that the model relies solely on the existing interaction network, which might vary across different tissue types [63, 64]. Although the network quality might not affect the model performance given the strong signal from gene expression information, the quality plays an important role in the interpretation step. Apart from the network quality, visualisation of a large network (>20,000 nodes) might be limited and hinder us from pinpointing small important subnetworks.

We propose Grandline by focusing on two key challenges of machine learning models for phenotype predictions [65], including the ability to model nonlinear relationships between data and outcomes and the incorporation of prior knowledge. Our GCN-based approach can be adapted for other applications. For example, instead of building a supervised model for predicting phenotypes, an autoencoder can be constructed for sample clustering based on protein interaction network and gene expression profile. In future work, other types of omics information such as mutation profile might be integrated as new channels to each node, allowing a model to learn comprehensive multi-omic patterns.

## Conclusions

We propose a deep graph convolutional neural network *Grandline* to integrate protein-protein interaction (PPI) network and gene expression to predict phenotypes, as well as identify biological subnetworks that are important for prediction. We demonstrate that Grandline correctly predicts phenotypes and identifies subnetworks that are relevant to each phenotype on simulated datasets, which capture both linear and nonlinear relationships between features values and target classes. In the nonlinear case, while some state-of-the-art models could achieve high predictive performance, only Grandline could pinpoint correct important subnetworks. Evaluations on real-world datasets, including major depressive disorder and cancer drug response, show that Grandline outperforms widely used machine learning models (ElasticNet logistic regression, SVM-Linear, SVM-RBF, Naïve Bayes and K-Nearest Neighbor). Compared to a standard deep neural network (DNN) model, Grandline could achieve similar prediction accuracy by requiring a lesser number of trainable parameters, allowing the model to be applied on a small-sized biological dataset. In terms of interpretability, Grandline is able to identify subnetworks in the PPI network (i.e., biological pathway) that are relevant to the underlying mechanisms of diseases and drug responses. Combining a deep learning model with graph structure and signal processing techniques has enabled us to capture complex relationships between gene expression patterns and phenotypes and integrate prior knowledge, which are the key challenges of applying machine learning in the fields of biology and medicine.

## Methods

### Data and data preprocessing

#### Real-world datasets

Gene expression profiles of healthy samples and patients with major depressive disorder (MDD) were obtained from two MDD studies [40, 41]. Drug response and gene expression profiles of 818 cancer cell lines and five drugs with the highest standard deviation of response values were obtained from the GDSC dataset [21]. The number of samples, genes, and phenotypic classes are described in **Table 1**. For MDD datasets, the target values were set to 0 for control and 1 for disease samples. For cancer drug response datasets, we set the target value to 1 when a cell line is sensitive to a given drug (i.e., the half-maximal inhibitory concentration, or IC50 value, is less than or equal to the maximum drug concentration used in the experiment) and 0 when a cell line is resistant to a given drug.

For gene expression profiles, TMM normalisation [5] was applied to the original raw read counts obtained from the studies, allowing the expression values to be comparable across genes and samples. We then calculated log2 fold-change with respect to the median expression of each gene to obtain the final normalised features for training and validating the models.

A protein-protein interaction network was obtained from STRING v10.5 [66], where high confidence interactions (edges with STRING score ≥800) were selected. In total, 17,181 nodes and 420,513 edges in the PPI network were selected. The specific numbers of nodes (genes) and edges (interactions) for each dataset were indicated in **Table 1**. We used networkx Python package [67] to process the graph and Gephi software [68] for graph visualisation. For pathway analysis, we used DAVID [10] to find KEGG pathways associated with important genes identified by Grandline. We also performed GSEA analysis to identify differentially expressed pathways using (gseapy; KEGG_2019_Human) [9].

#### Synthetic datasets

Synthetic data were used for evaluating if a model can correctly predict the target classes and identify correct predefined subnetworks that are related to the target classes. Two simulated datasets were generated to represent both linear (SIM-L) and nonlinear (SIM-NL) relationships between gene expression and binary target values. Each simulated dataset consists of 200 samples and 5,000 genes. Networkx package was used for generating a corresponding simulated graph of 5,000 nodes, where each node in the network represents a gene, and each edge represents protein-protein interaction. The simulated network is a random partition graph consists of 20 node clusters representing different gene clusters. Node signals (*X*) represent gene expression levels in a given sample, and target values (*y*) represent sample phenotypes.

For SIM-L, we defined *X* = *logit*(*y*)*w*^−1^, where *X* ∈ ℝ^200×5000^ contains a signal of 5000 nodes for 200 samples, *y* ∈ ℝ ^200^ is a target vector, and *w* ∈ ℝ^5000^ contains a predefined weight of each node. Since there are 20 node clusters, a similar weight was assigned to the nodes within the same cluster, resulting in 20 sets of node values associated with binary target values (**Supplementary figure 7A**). For SIM-NL, we generated a target vector *y* ∈ℝ^200^, signal matrix *X* ∈ ℝ^200×5000^ and *w* ∈ ℝ^5000^ that correspond to our predefined nonlinear relationships between the signal of nodes from 8 selected clusters and the target values. As a result, we obtained 200 sets of node values nonlinearly associated with binary target values (**Supplementary figure 7B**).

### Grandline framework

#### Graph convolutional network (GCN)

A PPI network is an undirected graph *G*(*V, E*), where |*V*| = *n* vertices (genes) and *E* is a set of edges (interaction). A graph signal *x* ∈ *R*^*n*^ contains node signals, i.e., gene expression values of each sample, where *x*_*i*_ is the value of *i*^*th*^ gene. Let *A* be an adjacency matrix and *D* be a diagonal matrix containing node degrees, the normalised graph Laplacian matrix is defined by

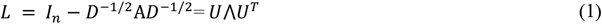

where *I*_*n*_ is an identity matrix, *U* is an eigenvector matrix and ⋀ is a diagonal matrix of eigenvalues. Similar to [32, 37], we defined a spectral graph convolution on *G* with a filter *g*_*θ*_ in the Fourier domain as

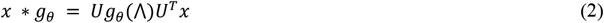

where *U*^*T*^*x* is the graph Fourier transform of *x, g*_*θ*_(⋀)= *diag*(*θ*) and *θ* ∈ *R*^*n*^ is a vector of Fourier coefficients.

To approximate the filter *g*_*θ*_, we used Chebychev polynomial degree *K* − 1 defined by

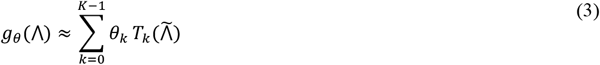

where *θ* ∈ *R*^*K*^ is a vector of Chebyshev coefficients, 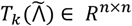 is the Chebyshev polynomial of order 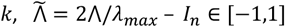 and *λ*_*max*_ is the largest eigenvalue of *L*. We note that the Chebyshev polynomials were calculated from *T*_*k*_(*x*)= 2*xT*_*k*−1_(*x*) − *T*_*k*−2_(*x*) with *T*_0_(*x*)= 1, *T*_1_(*x*)= *x* [38]. Thus, from Equation 1-3, the convolution of a graph signal *x* with filter *g*_*θ*_ is

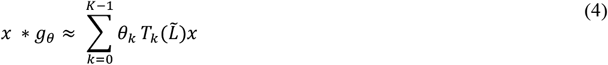

where 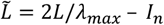.

To allow the pooling operation in the convolutional neural network, we followed a graph coarsening method implemented in [37]. Specifically, the PPI network was coarsened to create a multilevel partitioning of graph *G* by following Graclus’ greedy rule [69]. This results in a balanced binary tree, where the nodes were rearranged to facilitate a pooling step, and each pooling layer reduces the number of nodes by half according to the balanced binary tree.

Our GCN model was implemented by using Tensorflow v2.0 (with a custom GCN layer), binary cross-entropy as a loss function and Adam optimiser, followed the algorithm presented in [37]. The training process was controlled by a TensorFlow callback function, where the training is stopped when the validation accuracy was not improved (min_delta=0.001) for 15 epochs. Model configurations, including the number of layers, filters, batch size, as well as regularisation parameters, were systematically selected via cross-validation (**Table 2**; **Supplementary file 3**). Other utility functions such as graph coarsening were adapted from the existing Python script [37], where we modified the script to support a graph with multiple components.

#### Deciphering GCN via Grad-CAM

To identify subnetworks of the PPI network that are important for predicting a certain phenotype, we adopted Grad-CAM technique [36] to generate an importance heatmap on the input graph based on the feature map and the target classes. Originally, this technique is used for overlaying a heatmap onto the input image of a CNN model to highlight an important region of the image in predicting a given predicted class.

We defined *A*^*k*^ ∈ *G*(*V*_*l*_, *E*_*l*_) as a coarsen graph of a feature map *k* in last convolution layer and 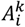 represents node *i*. Based on the predicted score of a given sample in class *c* (*Y*^*c*^), we calculated a gradient of *Y*^*c*^ with respect to the graph of feature map *k*,

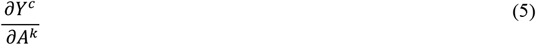

Next, we calculated the importance of feature map 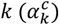 for the target class *c* by summing up the gradient of *Y*^*c*^,

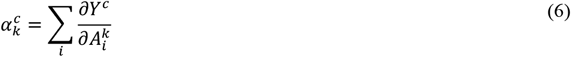

The node importance heatmap of the last convolution layer was calculated using

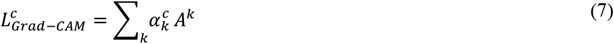

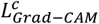 is the summation of gradients across all feature maps and captures importance value of each node in the coarsen graph.

To assign importance values to all nodes in the original graph *G*(*V,E*), we used the balanced binary tree from the coarsening step to infer importance values from each pooled node to the original nodes. To visualise important nodes in a graph, we calculated absolute importance values, sum up the values for each node/gene across multiple samples for each class, and highlighted the top genes (20% for SIM-L, 25% for SIM-NL, 5% for MDD and cancer drug response datasets, **Supplementary figure 8**).

### Related methods

We used scikit-learn [70] to train 5 state-of-the-art classification models (ElasticNet logistic regression, SVM-Linear, SVM-RBF, Naïve Bayes, and KNN). Multiple hyperparameters were tested and the best parameters were chosen as follow: C=[0.001, 0.01, 0.1, 1, 2, 5, 10] for ElasticNet and SVM models, gamma=[1e-6, 1e-5, 1e-4, ‘scale’] for SVM-RBF, var_smoothing=[1e-1, 1e-3, 1e-6, 1e-9, 1e-12, 1e-15] for Naïve Bayes, and k=[1, 5, 10, 15, 20, 25, 30, 35, 40, 45] for KNN. The baseline performance was based on predicting the majority class.

Beside the traditional machine learning models, multiple configurations of DNN models were evaluated using number of layers=[2,3], layer size=[16, 32, 64, 128, 256, 512], L2 regularization=[1e-1, 1e-2, 1e-3, 1e-4, 1e-5], lr=1e-4 and batch size=[16, 32, 64, …, n] where n is a number of samples. Similar to the GCN model, we used binary cross-entropy as a loss function, Adam optimizer, and the same early stopping criteria.

### Model training and evaluation

To measure the predictive performance, we held out 20% of the data for the final test. For the remaining 80%, we generated 10 random splits (80% for training and 20% for validation). MDD2 has a relatively small sample size, so all data were used for train and validation. We also enforced the test and validation set to have equal numbers of samples in each class, allowing us to validate the performance in an unbiased manner. The prediction accuracy, AUROC, AUPR were calculated based on the test set, and the average performance was reported (**Supplementary file 3**).

## List of abbreviations

DEGs: Differentially expressed genes
PPI: Protein-protein interaction
MDD: Major depressive disorder
DNN: Deep neural network
CNN: Convolutional neural network
GCN: Graph convolutional neural network
Grad-CAM: Gradient-weighted Class Activation Mapping
SIM-L: Simulated dataset with a linear relationship
SIM-NL: Simulated dataset with a nonlinear relationship
Linear-EN: ElasticNet model
KNN: K-Nearest Neighbor

## Supplementary Information

Supplementary file 1. Pathway enrichment results based on DAVID.

Supplementary file 2. GSEA results for major depressive disorder.

Supplementary file 3. Model performances.

## Declarations

### Ethics approval and consent to participate

Not applicable.

### Consent for publication

Not applicable.

### Availability of data and materials

Source code and example notebook for model training and prediction are available at www.github.com/BioML-CM/Grandline.

### Authors’ contributions

H.Y. developed Grandline and performed all computational analyses. C.S. planned and designed the project. H.Y. and C.S. wrote the manuscript.

## Acknowledgements

We thank Dr Niranjan Nagarajan for valuable insights and comments on the manuscript.

## References

1. Ding L, Bailey MH, Porta-Pardo E, Thorsson V, Colaprico A, Bertrand D, et al. Perspective on Oncogenic Processes at the End of the Beginning of Cancer Genomics. Cell. 2018;173:305-320.e10.

2. Lee J-K, Liu Z, Sa JK, Shin S, Wang J, Bordyuh M, et al. Pharmacogenomic landscape of patient-derived tumor cells informs precision oncology therapy. Nat Genet. 2018;50:1399–411.

3. Jansen R, Penninx Bwjh, Madar V, Xia K, Milaneschi Y, Hottenga JJ, et al. Gene expression in major depressive disorder. Mol Psychiatry. 2016;21:339–47.

4. Wang Z, Gerstein M, Snyder M. RNA-Seq: a revolutionary tool for transcriptomics. Nat Rev Genet. 2009;10:57–63.

5. Robinson MD, Oshlack A. A scaling normalization method for differential expression analysis of RNA-seq data. Genome Biol. 2010;11:R25.

6. Matthew J. Callow and TPSDSYHY. Statistical methods for identifying differentially expressed genes in replicated cDNA microarray experiments. Stat Sin. 2002;:111–39.

7. Robinson MD, McCarthy DJ, Smyth GK. edgeR: a Bioconductor package for differential expression analysis of digital gene expression data. Bioinformatics. 2009;26:139–40.

8. Love MI, Huber W, Anders S. Moderated estimation of fold change and dispersion for RNA-seq data with DESeq2. Genome Biol. 2014;15:1–21.

9. Subramanian A, Tamayo P, Mootha VK, Mukherjee S, Ebert BL, Gillette MA, et al. Gene set enrichment analysis: A knowledge-based approach for interpreting genome-wide expression profiles. Proc Natl Acad Sci U S A. 2005;102:15545–50.

10. Sherman BT, Lempicki RA, others. Systematic and integrative analysis of large gene lists using DAVID bioinformatics resources. Nat Protoc. 2009;4:44.

11. D’haeseleer P. How does gene expression clustering work? Nat Biotechnol. 2005;23:1499–501.

12. Hoadley KA, Yau C, Hinoue T, Wolf DM, Lazar AJ, Drill E, et al. Cell-of-Origin Patterns Dominate the Molecular Classification of 10,000 Tumors from 33 Types of Cancer. Cell. 2018;173:291-304.e6.

13. de Souto Mcp, Costa IG, de Araujo Dsa, Ludermir TB, Schliep A. Clustering cancer gene expression data: a comparative study. BMC Bioinformatics. 2008;9:497.

14. Adam G, Rampášek L, Safikhani Z, Smirnov P, Haibe-Kains B, Goldenberg A. Machine learning approaches to drug response prediction: challenges and recent progress. NPJ Precis Oncol. 2020;4:1–10.

15. Shipp MA, Ross KN, Tamayo P, Weng AP, Kutok JL, Aguiar RCT, et al. Diffuse large B-cell lymphoma outcome prediction by gene-expression profiling and supervised machine learning. Nat Med. 2002;8:68– 74.

16. Baptista D, Ferreira PG, Rocha M. Deep learning for drug response prediction in cancer. Brief Bioinform. 2020.

17. West M, Blanchette C, Dressman H, Huang E, Ishida S, Spang R, et al. Predicting the clinical status of human breast cancer by using gene expression profiles. Proc Natl Acad Sci U S A. 2001;98:11462–7.

18. Zhou J, Theesfeld CL, Yao K, Chen KM, Wong AK, Troyanskaya OG. Deep learning sequence-based ab initio prediction of variant effects on expression and disease risk. Nat Genet. 2018;50:1171–9.

19. Yeoh EJ, Ross ME, Shurtleff SA, Williams WK, Patel D, Mahfouz R, et al. Classification, subtype discovery, and prediction of outcome in pediatric acute lymphoblastic leukemia by gene expression profiling. Cancer Cell. 2002;1:133–43.

20. Orange DE, Agius P, DiCarlo EF, Robine N, Geiger H, Szymonifka J, et al. Identification of Three Rheumatoid Arthritis Disease Subtypes by Machine Learning Integration of Synovial Histologic Features and RNA Sequencing Data. Arthritis Rheumatol. 2018;70:690–701.

21. Iorio F, Knijnenburg TA, Vis DJ, Bignell GR, Menden MP, Schubert M, et al. A Landscape of Pharmacogenomic Interactions in Cancer. Cell. 2016;166:740–54.

22. Barretina J, Caponigro G, Stransky N, Venkatesan K, Margolin AA, Kim S, et al. The Cancer Cell Line Encyclopedia enables predictive modelling of anticancer drug sensitivity. Nature. 2012;483:603–7.

23. Zou H, Hastie T. Regularization and variable selection via the elastic net. J R Stat Soc Ser B Stat Methodol. 2005;67:301–20.

24. Suykens JAK, Vandewalle J. Least squares support vector machine classifiers. Neural Process Lett. 1999;9:293–300.

25. Cover TM, Hart PE. Nearest Neighbor Pattern Classification. IEEE Trans Inf Theory. 1967;13:21–7.

26. Szklarczyk D, Franceschini A, Wyder S, Forslund K, Heller D, Huerta-Cepas J, et al. STRING v10: Protein-protein interaction networks, integrated over the tree of life. Nucleic Acids Res. 2015;43:D447– 52.

27. Gawehn E, Hiss JA, Schneider G. Deep Learning in Drug Discovery. Molecular Informatics. 2016;35:3– 14.

28. Castelvecchi D. Can we open the black box of AI? Nat News. 2016;538:20.

29. Marcus G. Deep Learning: A Critical Appraisal. arXiv Prepr 180100631. 2018.

30. Krizhevsky A, Sutskever I, Hinton GE. ImageNet classification with deep convolutional neural networks. Commun ACM. 2017;60:84–90.

31. Xu L, Ren JS, Liu C, Jia J. Deep Convolutional Neural Network for Image Deconvolution. Adv Neural Inf Process Syst. 2014;27:1790–8.

32. Kipf TN, Welling M. Semi-Supervised Classification with Graph Convolutional Networks. 5th Int Conf Learn Represent ICLR 2017 - Conf Track Proc. 2016.

33. Nguyen T, Nguyen T, Le D-H. Graph convolutional networks for drug response prediction. bioRxiv. 2020;:2020.04.07.030908.

34. Oskooei A, Born J, Manica M, Subramanian V, Sáez-Rodríguez J, Martínez MR. PaccMann: Prediction of anticancer compound sensitivity with multi-modal attention-based neural networks. arXiv Prepr 181106802. 2018.

35. Yang J, Li A, Li Y, Guo X, Wang M. A novel approach for drug response prediction in cancer cell lines via network representation learning. Bioinformatics. 2019;35:1527–35.

36. Selvaraju RR, Cogswell M, Das A, Vedantam R, Parikh D, Batra D. Grad-CAM: Visual Explanations from Deep Networks via Gradient-based Localization. Proc IEEE Int Conf Comput Vis. 2017;:618–26.

37. Defferrard M, Bresson X, Vandergheynst P. Convolutional Neural Networks on Graphs with Fast Localized Spectral Filtering. Adv Neural Inf Process Syst. 2016;29:3844–52.

38. Mason, John C. and DCH. Chebyshev Polynomials. 2002.

39. Sundararajan M, Taly A, Yan Q. Axiomatic attribution for deep networks. In: International Conference on Machine Learning. 2017. p. 3319–28.

40. Le TT, Savitz J, Suzuki H, Misaki M, Teague TK, White BC, et al. Identification and replication of RNA-Seq gene network modules associated with depression severity. Transl Psychiatry. 2018;8:1–12.

41. Pantazatos SP, Huang YY, Rosoklija GB, Dwork AJ, Arango V, Mann JJ. Whole-transcriptome brain expression and exon-usage profiling in major depression and suicide: evidence for altered glial, endothelial and ATPase activity. Mol Psychiatry. 2017;22:760–73.

42. Gold PW, Licinio J, Pavlatou MG. Pathological parainflammation and endoplasmic reticulum stress in depression: potential translational targets through the CNS insulin, klotho and PPAR-$γ$ systems. Mol Psychiatry. 2013;18:154–65.

43. Yi Z, Li Z, Yu S, Yuan C, Hong W, Wang Z, et al. Blood-based gene expression profiles models for classification of subsyndromal symptomatic depression and major depressive disorder. PLoS One. 2012;7:e31283.

44. Su W-J, Zhang Y, Chen Y, Gong H, Lian Y-J, Peng W, et al. NLRP3 gene knockout blocks NF-$κ$B and MAPK signaling pathway in CUMS-induced depression mouse model. Behav Brain Res. 2017;322:1–8.

45. Grassi-Oliveira R, Brieztke E, Teixeira A, Pezzi JC, Zanini M, Lopes RP, et al. Peripheral chemokine levels in women with recurrent major depression with suicidal ideation. Brazilian J Psychiatry. 2012;34:71–5.

46. Bajetto A, Bonavia R, Barbero S, Schettini G. Characterization of chemokines and their receptors in the central nervous system: physiopathological implications. J Neurochem. 2002;82:1311–29.

47. Lall GS, Atkinson LA, Corlett SA, Broadbridge PJ, Bonsall DR. Circadian entrainment and its role in depression: a mechanistic review. J Neural Transm. 2012;119:1085–96.

48. Hasin DS, Goodwin RD, Stinson FS, Grant BF. Epidemiology of major depressive disorder: results from the National Epidemiologic Survey on Alcoholism and Related Conditions. Arch Gen Psychiatry. 2005;62:1097–106.

49. Van Hemert S, Hoekman AJW, Smits MA, Rebel JMJ. Immunological and gene expression responses to a Salmonella infection in the chicken intestine. Vet Res. 2007;38:51–63.

50. Bakeer MS, Youssef MI, Elshazly HM, Abdel-Samiee M, El-Gendy AA, Abouzed M, et al. On-treatment improvement of an emerging psychosomatic depressive disorder among salmonella carriers: a multicenter experience from Egypt. Infect Drug Resist. 2019;12:2573.

51. Karantza V. Keratins in health and cancer: More than mere epithelial cell markers. Oncogene. 2011;30:127–38.

52. Zhu L, Xiao F, Yu Y, Wang H, Fang M, Yang Y, et al. KSP inhibitor SB743921 inhibits growth and induces apoptosis of breast cancer cells by regulating p53, Bcl-2, and DTL. Anticancer Drugs. 2016;27:863–72.

53. Harada M, Ishihara Y, Itoh K, Yamanaka R. Kinesin superfamily protein-derived peptides with the ability to induce glioma-reactive cytotoxic T lymphocytes in human leukocyte antigen-A24+ glioma patients. Oncol Rep. 2007;17:629–36.

54. Cho RJ, Huang M, Campbell MJ, Dong H, Steinmetz L, Sapinoso L, et al. Transcriptional regulation and function during the human cell cycle. Nat Genet. 2001;27:48–54.

55. Yu MK, Ma J, Fisher J, Kreisberg JF, Raphael BJ, Ideker T. Visible Machine Learning for Biomedicine. Cell. 2018;173:1562–5.

56. Ma J, Yu MK, Fong S, Ono K, Sage E, Demchak B, et al. Using deep learning to model the hierarchical structure and function of a cell. Nat Methods. 2018;15:290.

57. Wu Z, Pan S, Chen F, Long G, Zhang C, Yu PS. A Comprehensive Survey on Graph Neural Networks. IEEE Trans Neural Networks Learn Syst. 2020;:1–21.

58. Dutil F, Cohen JP, Weiss M, Derevyanko G, Bengio Y. Towards Gene Expression Convolutions using Gene Interaction Graphs. arXiv Prepr 180606975. 2018.

59. Li Y, Kuwahara H, Yang P, Song L, Gao X. PGCN: Disease gene prioritization by disease and gene embedding through graph convolutional neural networks. bioRxiv. 2019;00:532226.

60. Han P, Yang P, Zhao P, Shang S, Liu Y, Zhou J, et al. GCN-MF: disease-gene association identification by graph convolutional networks and matrix factorization. In: Proceedings of the 25th ACM SIGKDD international conference on knowledge discovery & data mining. 2019. p. 705–13.

61. Ying R, Bourgeois D, You J, Zitnik M, Leskovec J. Gnnexplainer: Generating explanations for graph neural networks. Adv Neural Inf Process Syst. 2019;32:9240.

62. Schnake T, Eberle O, Lederer J, Nakajima S, Schütt KT, Müller KR, et al. Higher-order explanations of graph neural networks via relevant walks. 200603589. 2020.

63. Bossi A, Lehner B. Tissue specificity and the human protein interaction network. Mol Syst Biol. 2009;5:260.

64. Magger O, Waldman YY, Ruppin E, Sharan R. Enhancing the Prioritization of Disease-Causing Genes through Tissue Specific Protein Interaction Networks. PLoS Comput Biol. 2012;8:e1002690.

65. Costello JC, Heiser LM, Georgii E, Gönen M, Menden MP, Wang NJ, et al. A community effort to assess and improve drug sensitivity prediction algorithms. Nat Biotechnol. 2014;32:1–103.

66. Szklarczyk D, Morris JH, Cook H, Kuhn M, Wyder S, Simonovic M, et al. The STRING database in 2017: quality-controlled protein–protein association networks, made broadly accessible. Nucleic Acids Res. 2017;45:D362–8.

67. Hagberg A, Swart P, Chult D. Exploring network structure, dynamics, and function using NetworkX. Proc 7th Python Sci Conf. 2008;:11–15.

68. Bastian M;, Heymann S;, Jacomy M. Gephi : An Open Source Software for Exploring and Manipulating Networks. Icwsm. 2009;8:361–2.

69. Dhillon IS, Guan Y, Kulis B. Weighted graph cuts without eigenvectors a multilevel approach. IEEE Trans Pattern Anal Mach Intell. 2007;29:1944–57.

70. Pedregosa F, Varoquaux G, Gramfort A. scikit-learn: Machine Learning in Python. Journal of Machine Learning Research. 2011;:2825–30. http://scikit-learn.org/.

