## Supplementary figures for "Network-guided supervised learning on gene expression using a graph convolutional neural network"

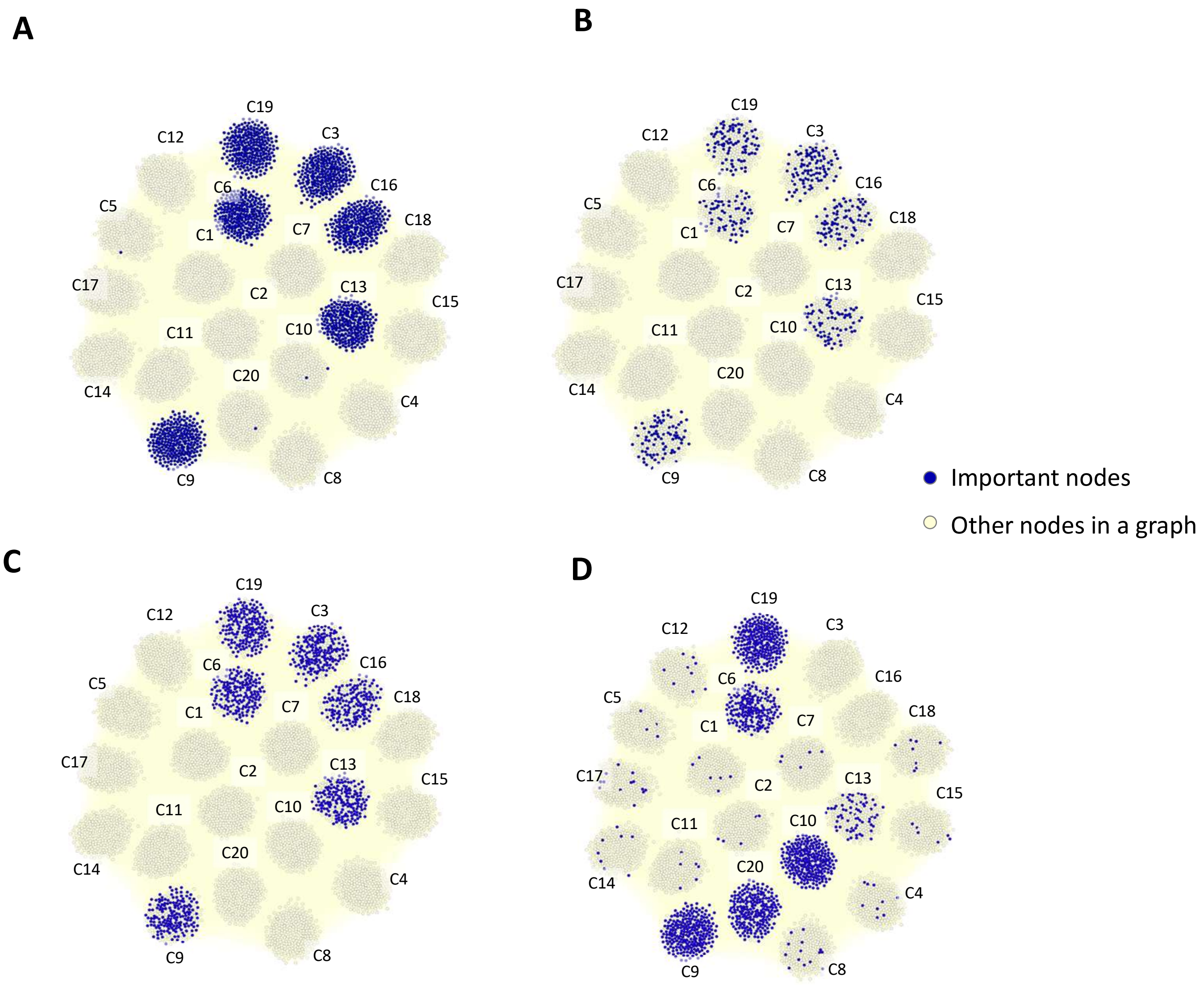


**Supplementary figure 1.** Important nodes identified by SVM-Linear (A) and Linear-EN (B) for synthetic dataset with linear relationship between features and target values (SIM-L). (C) and (D) are the important features in DNN models for SIM-L (correct subnetworks include C3, 6, 9, 13, 16, 19) and SIM-NL (correct subnetworks include C3, 6, 9, 10, 13, 16, 19, 20) datasets, respectively. The importance value of each gene was calculated by summing up the importance obtained from the integrated gradients (IG) method. For SIM-NL, IG method could not pinpoint C3 and C16.

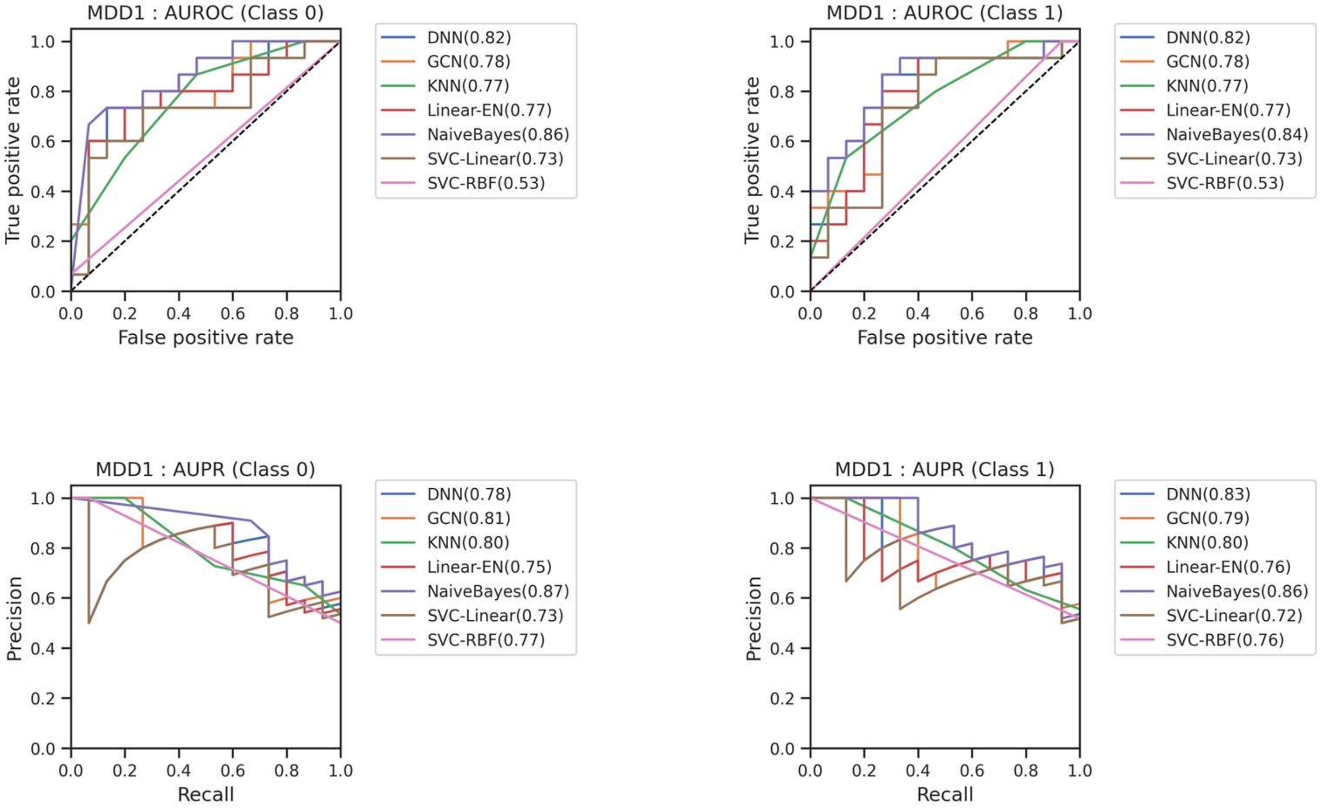


**Supplementary figure 2.** Area under a receiver operating characteristic (AUROC) and Area under the precision-recall curve AUPR for MDD1.

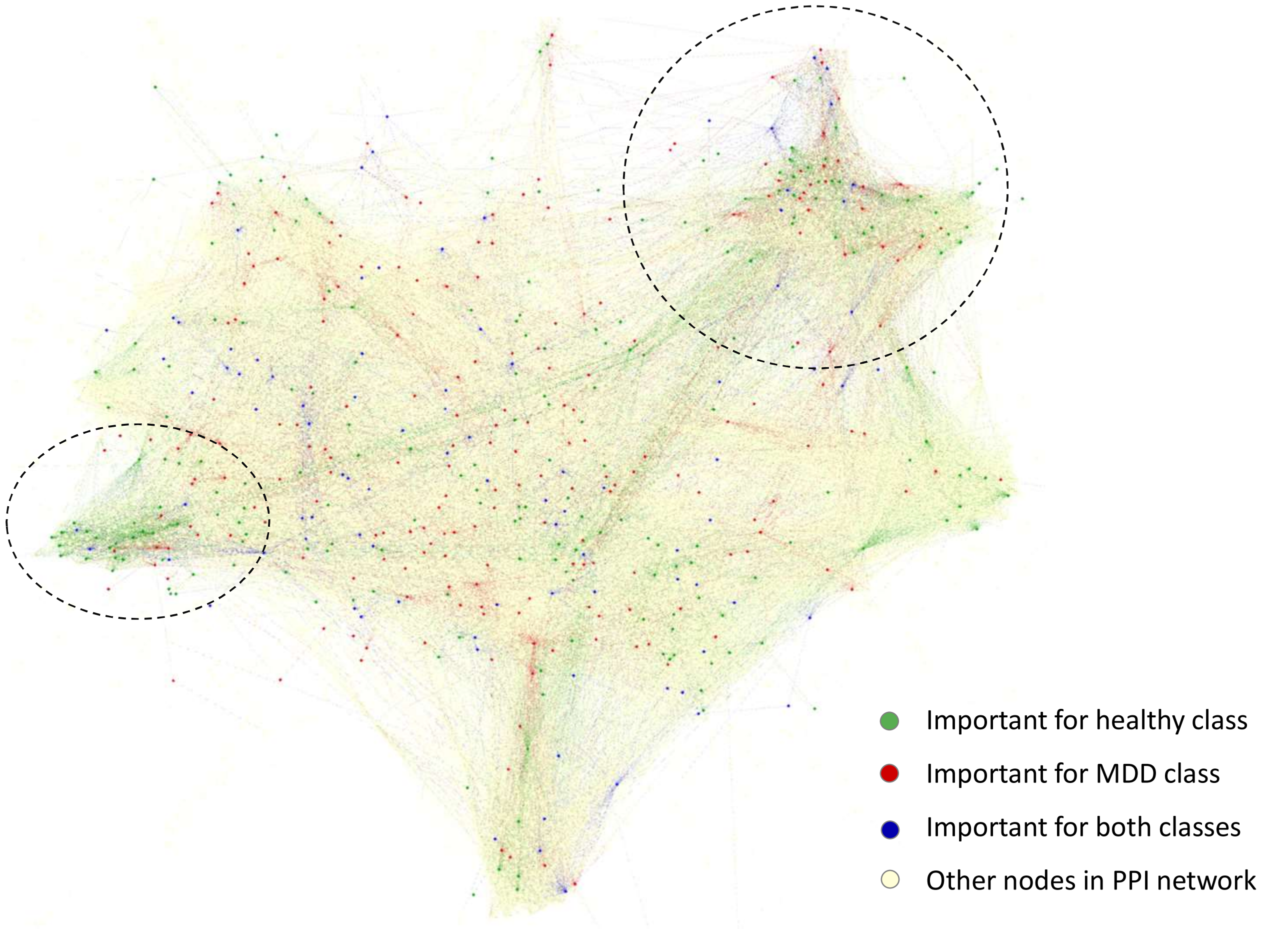


**Supplementary figure 3.** Important genes identified by Grandline for MDD1 dataset, highlighting subnetworks of PPI network that could distinguish healthy and MDD samples.

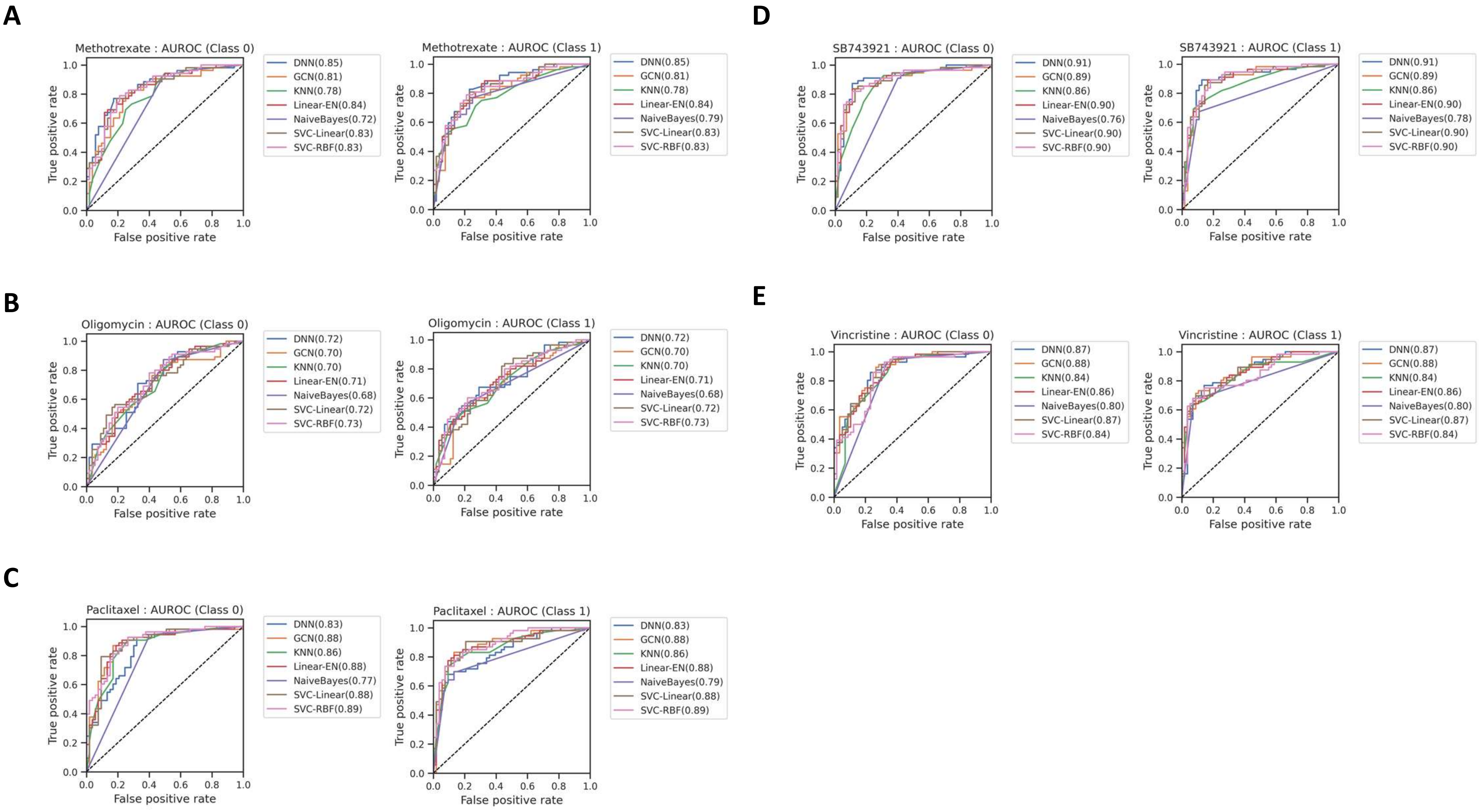


**Supplementary figure 4.** Area under a receiver operating characteristic (AUROC) for predicting cancer drug response

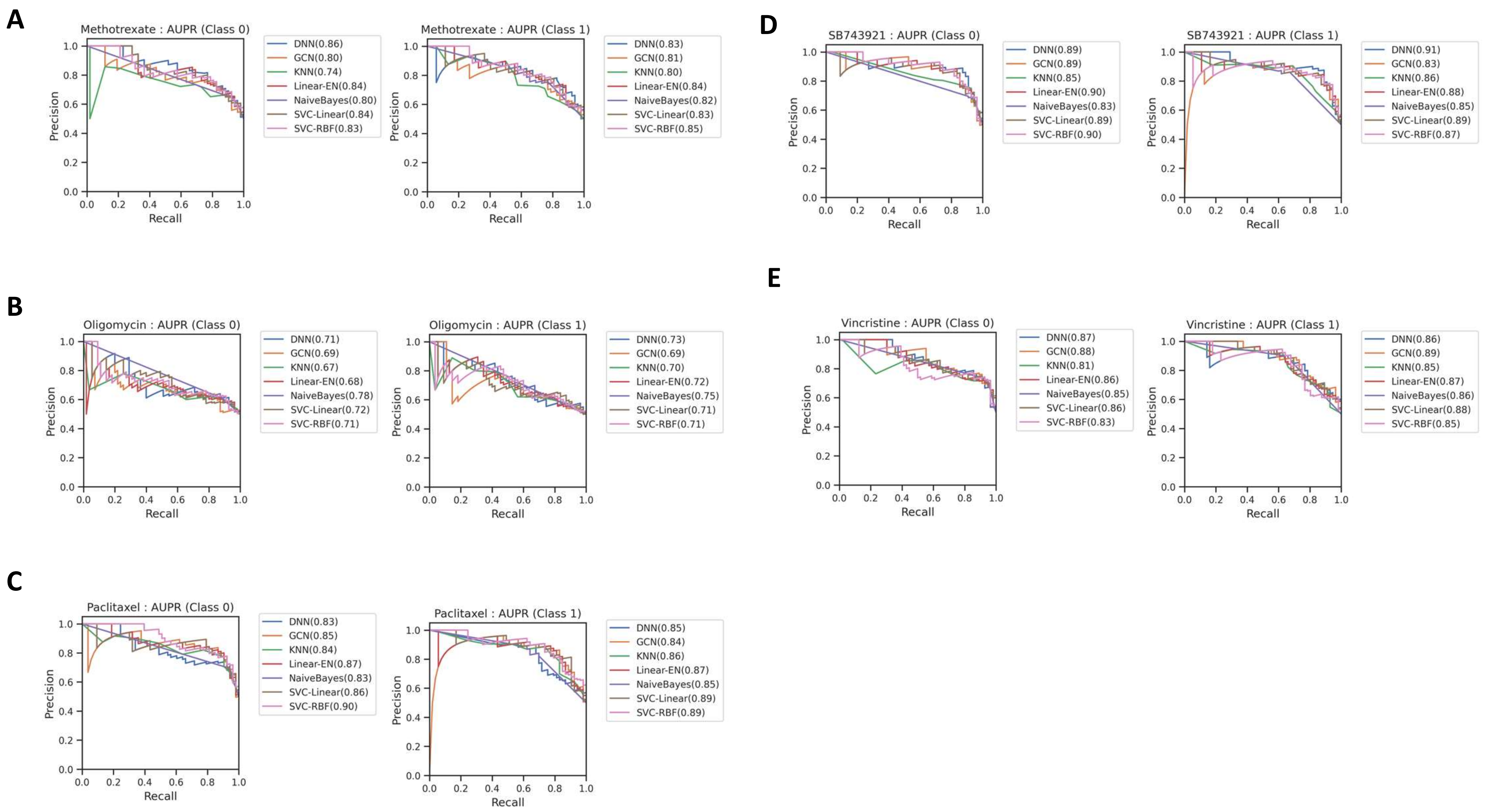


**Supplementary figure 5.** Area under the precision-recall curve (AUPR) for predicting cancer drug response

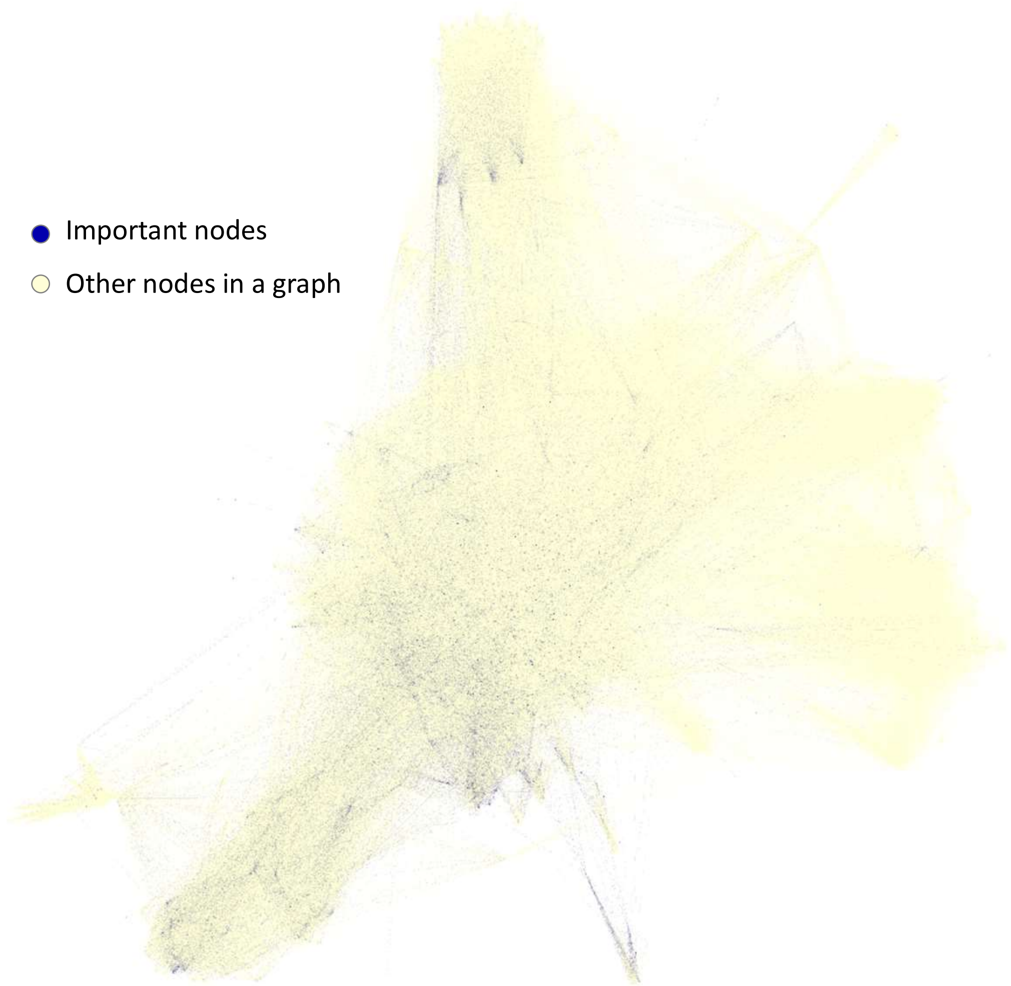


**Supplementary figure 6** Importance subnetwork identified by the integrated gradients methods on DNN model for SB743921

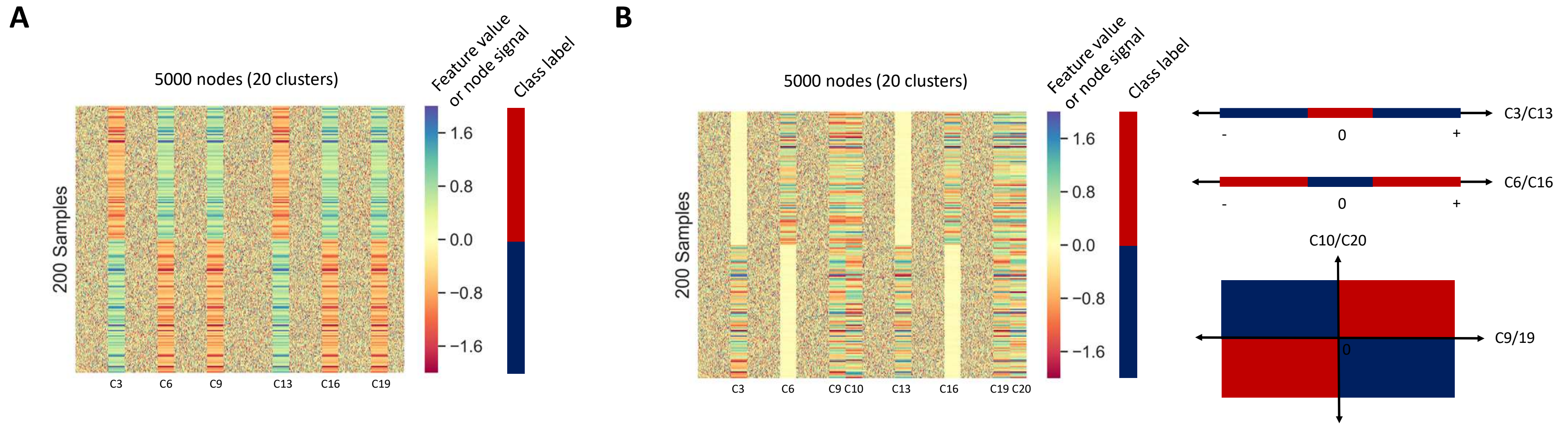


**Supplementary figure 7. Generating simulated datasets.** A random graph with 5000 nodes was generated with 20 clusters, i.e., sets of highly connected nodes. Next, 200 samples (100 samples for each class) were generated by assigning feature values of nodes in specific clusters to be linearly (A) and nonlinearly (B) associated with the class label, while feature values of the remaining nodes were assigned randomly.

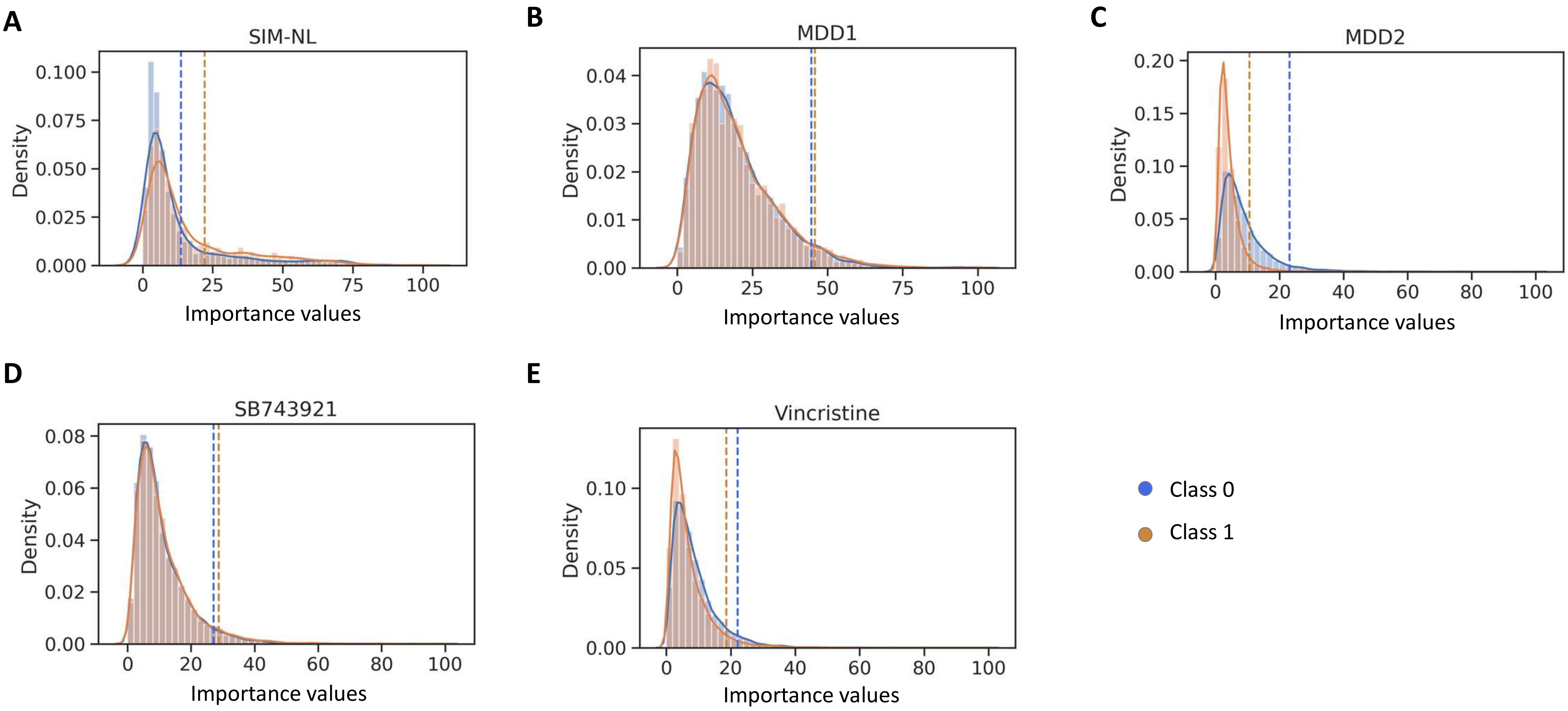


**Supplementary figure 8.** Histogram of the normalised absolute importance values for each class. The dashed lines represent the thresholds used for identifying important nodes.
